# Engineering isoprenoid quinone production in yeast

**DOI:** 10.1101/2020.02.06.932020

**Authors:** Divjot Kaur, Christophe Corre, Fabrizio Alberti

## Abstract

Isoprenoid quinones are bioactive molecules that include an isoprenoid chain and a quinone head. They are traditionally found to be involved in primary metabolism, where they act as electron transporters, but specialized isoprenoid quinones are also produced by all domains of life. Here, we report the engineering of a baker’s yeast strain, *Saccharomyces cerevisiae* EPYFA3, for the production of isoprenoid quinones. Our yeast strain was developed through overexpression of the shikimate pathway in a well-established recipient strain (*S. cerevisiae* EPY300) where the mevalonate pathway is overexpressed. As a proof of concept, our new host strain was used to overproduce the endogenous isoprenoid quinone coenzyme Q_6_, resulting in a final four-fold production increase. EPYFA3 represents a valuable platform for the heterologous production of high value isoprenoid quinones. EPYFA3 will also facilitate the elucidation of isoprenoid quinone biosynthetic pathways.

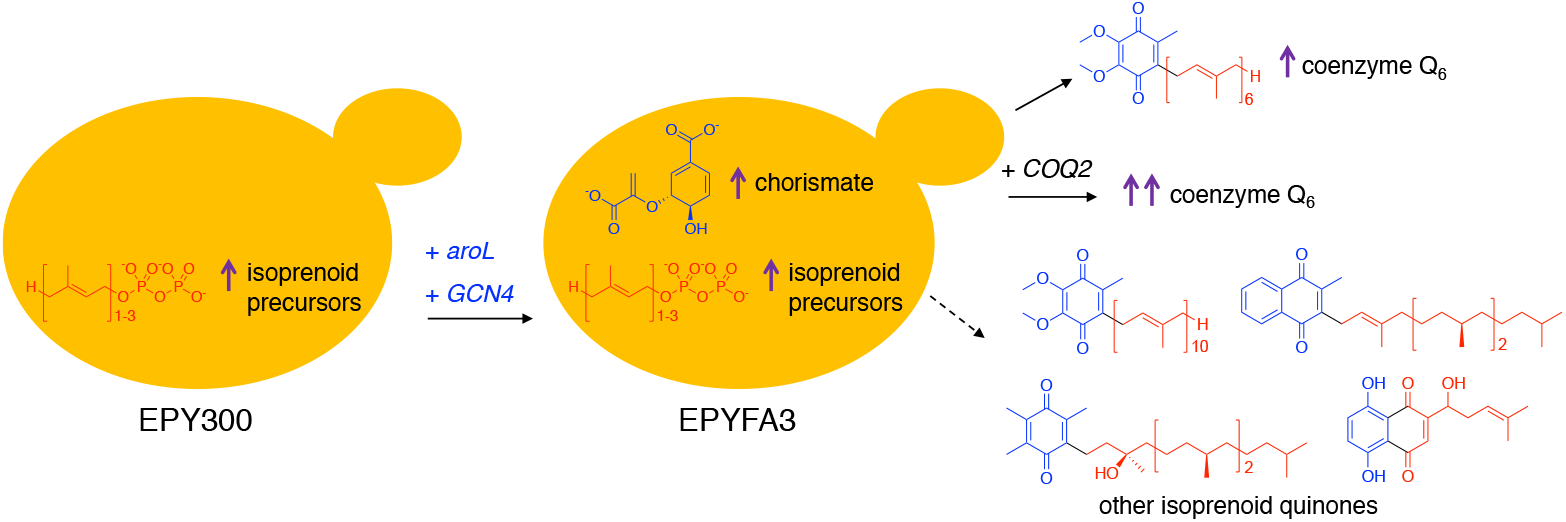

## INTRODUCTION

Isoprenoid quinones (IQs) are biologically active metabolites that are made of a hydrophobic isoprenoid tail and a polar quinone head.^1^ Traditionally, IQs have been found to act as electron transporters in respiratory and photosynthetic electron transport chains.^2^ In this context, the isoprenoid moiety allows IQs to be anchored to phospholipid bilayer membranes, whereas the polar head enables their interaction with proteins. Naphthoquinones (menaquinone and phylloquinone) contain a naphthalene-related quinone head, whereas benzoquinones (ubiquinone and plastoquinone) include a benzene-related quinone ring. IQs have mixed biosynthetic origins. Their isoprenoid chain comes from isopentenyl pyrophosphate (IPP) and dimethylallyl pyrophosphate (DMAPP).^3^ The quinone head can come from different sources, in particular from the *O*-succinylbenzoate pathway *via* chorismate or from the shikimate pathway *via* chorismate, tyrosine, or phenylalanine.^2,4^ Specialized IQs often include a polyketide head group, as commonly seen for fungal meroterpenoids.^5^ Ubiquinone-10 (coenzyme Q_10_),^1^ phylloquinone (vitamin K1)^6^ and α-tocopherol quinone (vitamin E quinone)^7^ are examples of widespread IQs (Figure 1). Species-specific IQs, which are generally regarded as specialized metabolites, include the antitumor furaquinocin A,^8^ the antiviral 4-hydroxypleurogrisein,^9^ the antioxidant naphterpins^10^ and the antitumor antibiotics BE-40644^11^ and shikonin (Figure 1).^12,13^

**Figure 1.**
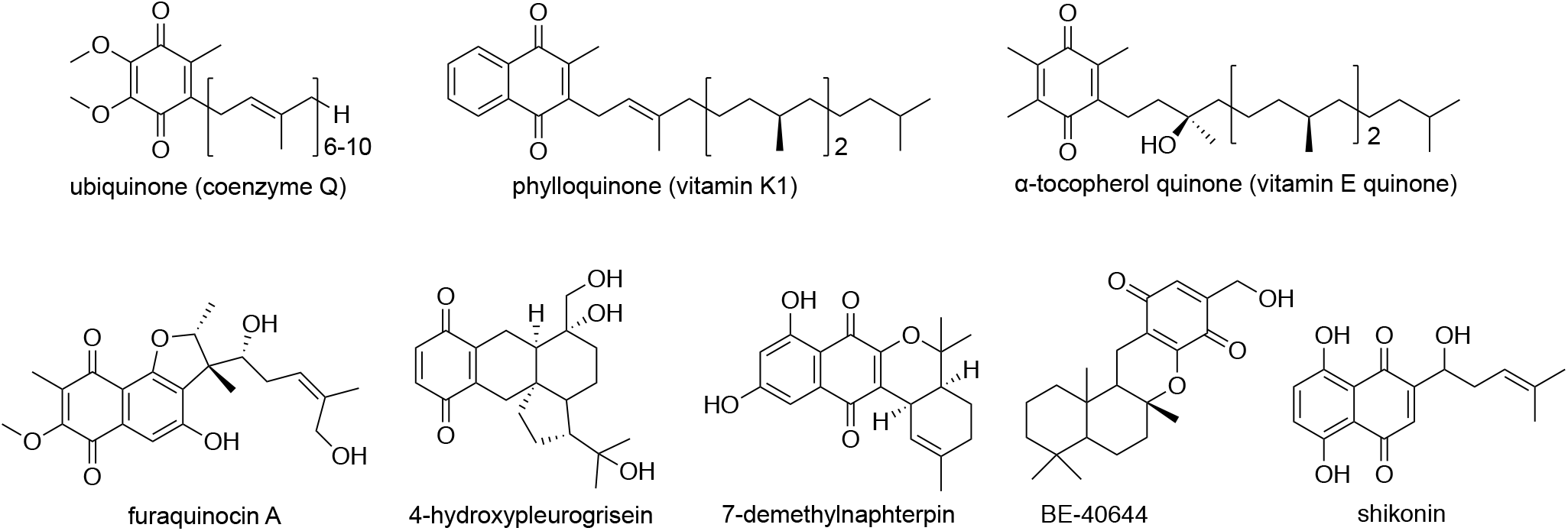
Examples of isoprenoid quinones.

The biosynthetic pathways to ubiquitous IQs have been investigated extensively, although a complete picture is still lacking for compounds such as coenzyme Q.^14^ The biosynthesis of specialized IQs is often directed by cryptic metabolic pathways. Their elucidation requires the use of interdisciplinary approaches, such as total synthesis and reconstitution of *in vitro* reactions with recombinant biosynthetic enzymes, as shown for instance by Murray *et al.*^10^ for the biosynthesis of naphterpins and marinones. To increase IQs production and to facilitate the elucidation of their cryptic biosynthetic routes, a suitable biological platform providing ample precursor supply is needed.

Here we report the engineering of a *Saccharomyces cerevisiae* (baker’s yeast) strain that overproduces the building blocks required for IQ biosynthesis.

As a proof of concept, we used our strain to improve the production of coenzyme Q_6_. *S. cerevisiae* has been metabolically engineered in the past to overproduce isoprenoids.^15^ Notably, the Keasling lab has engineered the mevalonate pathway in yeast, in order to provide increased amounts of IPP and DMAPP for the assembly of various downstream products, such as the antimalarial drug precursor artemisinic acid,^16^ the biofuel precursor bisabolene,^17^ as well as natural and unnatural cannabinoids.^18^ Strain *S. cerevisiae* EPY300 (Supporting Table S1) has been used to produce terpenoids, such as artemisinic acid^16^ and costunolide,^19^ in high amounts.

EPY300 includes two copies of a truncated, soluble variant of 3-hydroxy-3-methyl-glutaryl-coenzyme A reductase (*tHMGR*), as well as an additional copy of the farnesyl-pyrophosphate (FPP) synthase gene *ERG20*, a copy of the semi-dominant allele *upc2-1* for upregulation of genes involved in the biosynthesis of sterols, and a methionine inducible downregulation mechanism for the control of *ERG9*, which codes for squalene synthase (Scheme 1). In the present work, we acquired *S. cerevisiae* EPY300 from the Keasling lab and engineered it further to overexpress the shikimate pathway. The gene *aroL* codes for the shikimate kinase II in *Escherichia coli*,^20^ and its overexpression in *S. cerevisiae* has been shown to improve production of the shikimate pathway-derived *p*-coumaric acid.^21^ *GCN4* codes for a leucine zipper transcriptional activator (Gcn4p) in yeast, which is induced under amino acid starvation, and in turn triggers the upregulation of biosynthetic genes involved in the production of most amino acids.^22^ The targets of Gcn4p include genes *ARO1*, *ARO2*, *ARO3* and *ARO4*, which in yeast code for the enzymes that catalyze the seven steps of the shikimate pathway leading to chorismate. Chorismate can then be used for downstream processes, such as the biosynthesis of aromatic amino acids and that of the head group of IQs. Gcn4p also targets vitamin-cofactor biosynthetic genes, such as those that code for the enzymes responsible for the biosynthesis of NAD+, FAD and coenzyme-A, as well as for the biosynthesis of tryptophan, a precursor of nicotinamide, which in turn is a component of NAD+ and NADP+.^23^ Therefore, overexpression of *GCN4* can also lead to increased amounts of the cofactors that are required by secondary metabolite biosynthetic enzymes, including those oxidoreductases that catalyze the assembly of IQs. Eukaryotic cytochrome P450 enzymes, such as CYP76B74 involved in the biosynthesis of the IQ shikonin,^24^ require the cofactor NADPH as a source of electrons, and some dehydrogenases use NAD+ or NADP+ as electron acceptors.

## RESULTS AND DISCUSSION

In order to produce a yeast strain that would accumulate precursors to IQs in high amounts, we set out to overexpress the shikimate pathway in EPY300. To introduce the *E. coli aroL* and overexpress the endogenous *GCN4*, we assembled the multigene vector pFA011 using the MoClo Yeast Toolkit^25^ (see Methods for details on plasmid assembly). The sequence of the shikimate kinase II gene *aroL* (NCBI Gene ID: 945031) was codon-optimized for expression in *S. cerevisiae* (Supporting Table S2), whereas *GCN4* (NCBI Gene ID: 856709) was amplified from the genomic DNA of *S. cerevisiae* EPY300. The strong constitutive promoters *pTEF1* and *pTEF2* were placed upstream of *aroL* and *GCN4*, respectively, as they are known to drive steady gene expression in media with different carbon sources.^26^ Upon linearization with NotI, pFA011 was introduced into *S. cerevisiae* EPY300 *via* the LiAc/single-stranded carrier DNA/PEG method,^27^ producing strain EPYFA3 (Scheme 1). Correct integration in the *LEU2* locus was verified through PCR amplification of the *LEU2* 5’ region (Supporting Figure S3a), as well as of the *LEU2* 3’ region (Supporting Figure S3b). In order to confirm that the genes of the shikimate pathway were upregulated by the transcriptional activator Gcn4p in EPYFA3, we performed RT-qPCR analysis, targeting the aromatic amino acid biosynthetic genes *ARO1*, *ARO2*, *ARO3* and *ARO4*. The expression of *GCN4* itself was also evaluated in EPYFA3 and compared to that of the *GCN4* single copy-containing strain EPY300. Three independent colonies of each EPYFA3 and EPY300 were grown in synthetic defined (SD) broth with dextrose, shaking at 200 rpm, 30°C, for 24 hours. The pre-cultures were used to inoculate production cultures in SD broth with 0.1% dextrose, 2% galactose and 1 mM methionine, which were shaken at 200 rpm, 30°C, for 120 hours.

**Scheme 1.**
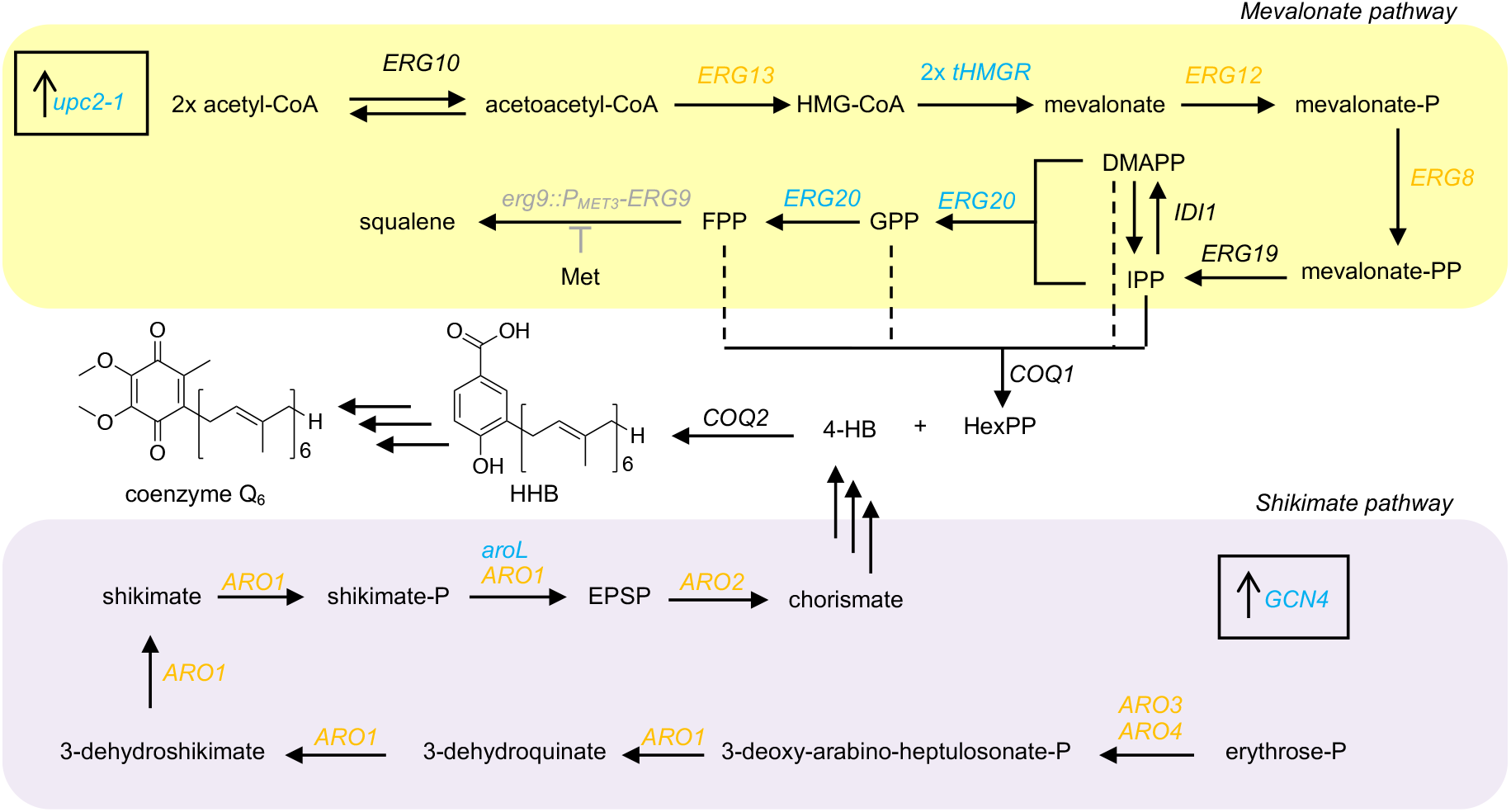
Schematic representation of the engineered metabolism of *S. cerevisiae* EPYFA3. Genes from the mevalonate pathway are grouped in the yellow panel (adapted from Ro *et al.*^16^); genes from the shikimate pathway are grouped in the purple panel. Overexpressed genes are highlighted in light blue; genes indirectly upregulated (from *upc2-1* for the mevalonate pathway and from *GCN4* for the shikimate pathway) are highlighted in orange. Repression of *ERG9* through the methionine-repressible promoter P_*MET3*_ is represented in grey. Usage of DMAPP, GPP or FPP by *COQ1* is shown through dashed lines as it is unclear which of these allylic pyrophosphates is used *in vivo* to make HexPP. The first reaction in the biosynthetic pathway to coenzyme Q_6_, the condensation of HexPP and 4-HB catalyzed by *COQ2*, is also shown. HMG-CoA: 3-hydroxy-3-methyl-glutaryl-coenzyme A; IPP: isopentenyl pyrophosphate; DMAPP: dimethylallyl pyrophosphate; GPP: geranyl pyrophosphate; FPP farnesyl pyrophosphate; HexPP: hexaprenyl pyrophosphate; 4-HB: 4-hydroxybenzoic acid; HHB: 3-hexaprenyl 4-hydroxybenzoic acid; EPSP: 5-enolpyruvylshikimate-3-phosphate.

Cells were harvested, RNA purified, converted into first-strand cDNA and used for RT-qPCR. The expression of *GCN4* was confirmed to be higher in EPYFA3 than in EPY300 by almost a five-fold (Figure 2), due to the introduction of a second copy of *GCN4* under control of the strong constitutive *pTEF2* promoter in EPYFA3. Upregulation of *ARO1*, *ARO2*, *ARO3* and *ARO4* was confirmed in EPYFA3 upon overexpression of *GCN4*, in accordance with what described by Natarajan *et al.*^22^ *ARO1* showed a 5.60 (± 1.00)-fold increase, *ARO2* a 2.86 (± 0.92)-fold increase, *ARO3* a 10.27 (± 1.40)-fold increase, and *ARO4* a 5.26 (± 0.82)-fold increase in EPYFA3 compared to EPY300 (Figure 2).

**Figure 2.**
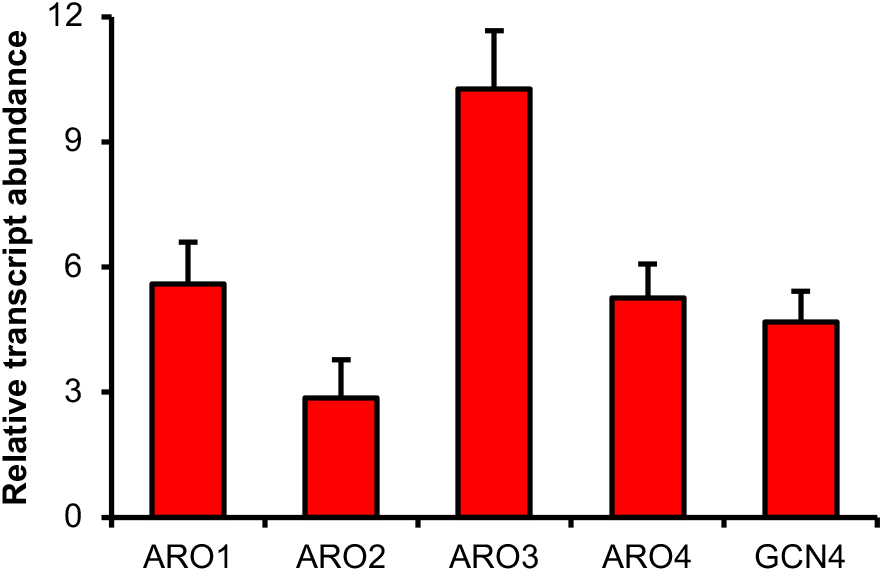
RT-qPCR expression analysis. Relative transcript abundance of genes *ARO1*, *ARO2*, *ARO3*, *ARO4* and *GCN4* in *S. cerevisiae* EPYFA3. Actin (*ACT1*) was used as the reference gene, and the average Ct value of each target gene in *S. cerevisiae* EPY300 was used as the calibrator (the expression of each gene in *S. cerevisiae* EPYFA3 is shown relative to the expression of the same gene in *S. cerevisiae* EPY300). Values and error bars reflect mean and standard deviation of three biological replicates.

Next, we tested the ability of *S. cerevisiae* EPYFA3 to be used as a host for the increased production of IQs. As a proof of concept, we decided to test for the overproduction of ubiquinone-6 (coenzyme Q_6_), since this is the yeast endogenous benzoquinone and is assembled by condensing building blocks deriving from the mevalonate and the shikimate pathways (see the review of Stefely and Pagliarini^14^ for an extensive overview of coenzyme Q biosynthesis). The quinone head of coenzyme Q_6_ in yeast comes primarily from 4-hydroxybenzoate (4-HB), as well as from 4-aminobenzoate through a secondary pathway that is only active in the absence of 4-HB.^28^ In turn, 4-HB either comes from tyrosine or is made *de novo* through the shikimate pathway. It should be noted that the overexpression of *GCN4* does not upregulate the tyrosine biosynthetic gene *TYR1*.^22^

Conversely, the isoprenoid tail of coenzyme Q_6_ in yeast comes from the mevalonate pathway, presumably as a result of the condensation of IPP with the allylic pyrophosphates DMAPP, geranyl-pyrophosphate (GPP) or FPP.^3^ To assess the ability to redirect building blocks towards a specific metabolite of interest in EPYFA3, we overexpressed the gene *COQ2*, which codes for the 4-HB polyprenyltransferase, a UbiA family prenyltransferase that catalyzes the first committed reaction in the biosynthesis of coenzyme Q_6_.^29^ The sequence of gene *COQ2* (NCBI Gene ID: 855778) was optimized to remove a BsmBI site through the introduction of a synonymous substitution (G885A). We assembled an expression vector, pFA004, that contained a copy of *COQ2* under control of the strong constitutive promoter *pCCW12* (see Methods for details on plasmid assembly). Upon linearization with NotI, pFA004 was introduced into both *S. cerevisiae* EPY300 and EPYFA3 *via* the LiAc/single-stranded carrier DNA/PEG method,^27^ producing strains EPYFA4 and EPYFA7 respectively. Correct integration in the *URA3* locus was verified through PCR amplification of the *URA3* 5’ region (Supporting Figure S3c), as well as of the *URA3* 3’ region (Supporting Figure S3d).

We determined the amount of coenzyme Q_6_ accumulated in strains EPY300, EPYFA3, EPYFA4 and EPYFA7. For each strain, three independent colonies were used to inoculate a pre-culture in SD broth with dextrose, shaking at 200 rpm, 30°C, for 24 hours. The pre-cultures were used to inoculate production cultures in SD broth with 0.1% dextrose, 2% galactose and 1 mM methionine, which were shaken at 200 rpm, 30°C, for 120 hours. Lipids were extracted from pelleted cells using a modified protocol from Marbois *et al.*^28^ (see Supporting Methods) and analyzed through reversed-phase HPLC diode-array base-peak chromatogram obtained at 274 nm. Quantification was performed using a coenzyme Q_6_ (Sigma-Aldrich) external standard and a calibration curve, through the integration of the area underneath the UV peak corresponding to coenzyme Q_6_ (Figure 3a,b). The identity of coenzyme Q_6_ in the yeast extracts was also confirmed through mass spectrometry (Supporting Figure S4). An increase in coenzyme Q_6_ production could be observed upon overexpression of the shikimate pathway in strain EPYFA3 (Figure 3c), which accumulated 17.2 (± 7.5) mg L^−1^, compared to EPY300, which produced 6.8 (± 0.4) mg L^−1^. Similarly, increased production was detected when *COQ2* was being overexpressed in EPYFA7, corresponding to 27.8 (± 8.7) mg L^−1^, compared to strain EPYFA4 where *COQ2* was also overexpressed but the shikimate pathway was not, which accumulated 18.7 (± 4.2) mg L^−1^. Interestingly, it could also be observed that the overexpression of the prenyl-transferase *COQ2* led to increased accumulation of coenzyme Q_6_ by comparing the titers obtained from strain EPYFA7 and EPYFA3, suggesting that the overexpression of a prenyltransferase biosynthetic gene for an IQ in EPYFA3 can successfully redirect building blocks from mevalonate and shikimate pathways to the assembly of a specific metabolite of interest.

**Figure 3.**
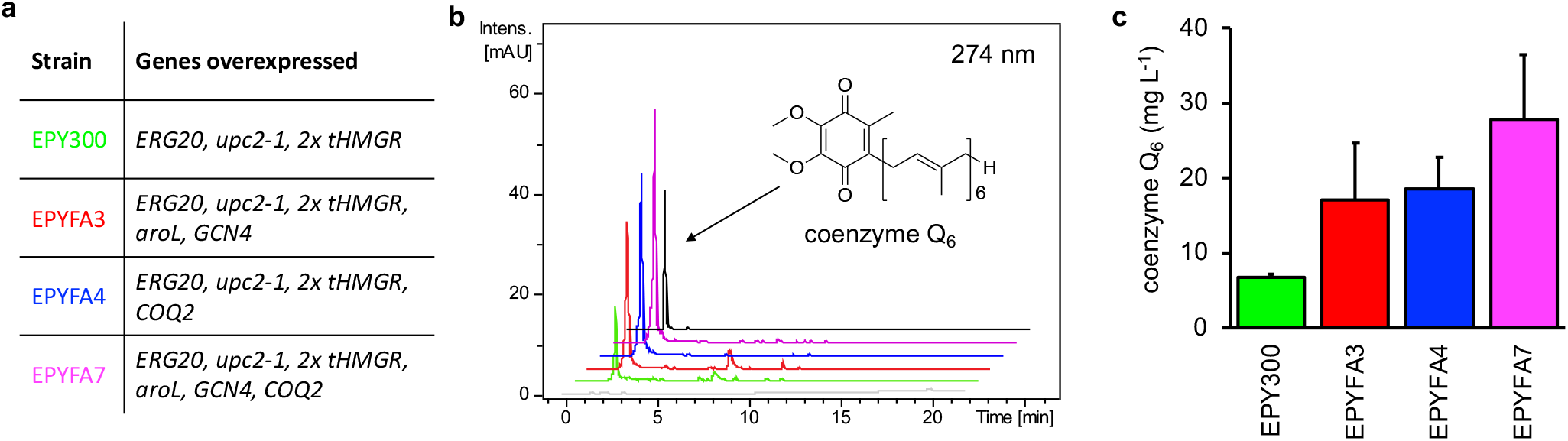
Quantification of coenzyme Q_6_ production. (a) List of *S. cerevisiae* strains with corresponding overexpressed genes. (b) HPLC diode-array base-peak chromatograms (274 nm) for detection of coenzyme Q_6_. Chromatograms from bottom to top: growth medium control (grey), EPY300 (green), EPYFA3 (red), EPYFA4 (blue), EPYFA7 (magenta), authentic standard (black). (c) Quantification of coenzyme Q_6_ from the four *S. cerevisiae* strains with HPLC-diode-array detection. Values and error bars reflect mean and standard deviation of three biological replicates.

It should be noted that methionine is used to downregulate the expression of the squalene synthase gene *ERG9* but it is also consumed during coenzyme Q_6_ biosynthesis in *S. cerevisiae*, as *S*-adenosylmethionine (SAM) is employed as a cofactor by the *O*-methyltransferase Coq3p in the catalysis of two coenzyme Q_6_ biosynthetic steps.^30^ However, the amount of coenzyme Q_6_ produced by EPYFA7 corresponds to 0.047 (± 0.015) mM. Therefore, only 0.094 (± 0.03) mM equivalent of methionine is used for the production of coenzyme Q_6_ by the highest producing strain, meaning that ample methionine is available to direct downregulation of *ERG9* through the methionine-repressible promoter P_*MET3*_ (Scheme 1).

The growth of the four yeast strains in shake-flask cultures was monitored measuring the OD_600_ at various timepoints (Supporting Figure S5). The growth of EPYFA3 and EPYFA7 appeared to be slower than their counterpart strains EPY300 and EPYFA4, showing a possible metabolic impairment due to the shikimate pathway being upregulated. Overexpression of the prenyltransferase gene *COQ2* appeared instead to increase cell growth, as it could be seen in strains EPYFA4 and EPYFA7 compared to EPY300 and EPYFA3, respectively, putatively due to an improved fitness upon accumulation of additional coenzyme Q_6_ or to a reduced toxicity of shikimate-derived pathway intermediates.

Noteworthily, since *S. cerevisiae* EPYFA3 is a derivative of BY4742, it harbors unused auxotrophic markers that can be employed to introduce other expression vectors. Loci *LYS2* (mutated to *lys2Δ0*), *URA3* (mutated to *ura3Δ0*) and *HO* can be used for the chromosomal integration of up to three plasmids. *LYS2*, *URA3*, as well as expression cassettes for resistance to antifungals, such as HygromycinR and KanMXR, can be used as selectable markers for the maintenance of integrative and/or replicative plasmids. This will allow EPYFA3 to be used to reconstitute the biosynthesis of IQs, as well as that of any other compounds derived from the mevalonate and the shikimate metabolic pathways.

In conclusion, we have engineered a yeast strain that overexpresses both the mevalonate and the shikimate pathways and can therefore be used to produce IQs in which the quinone ring derives from the shikimate pathway. We envisage that our strain will be particularly useful to express prenyl-transferase enzymes and other IQ biosynthetic enzymes, which have recently gained attention as potential biocatalysts.^31^ It will also allow to support accumulation of metabolites in amounts sufficient for purification and structure determination. Therefore, it will be a valuable platform to refactor and characterize the cryptic biosynthesis of specialized IQs for which a fast-growing heterologous host is needed.

## METHODS

### Reagents

All chemicals were purchased from Sigma-Aldrich, unless otherwise stated. Q5 High-Fidelity 2X Master Mix, restriction endonucleases, shrimp alkaline phosphatase (rSAP), and T4 DNA ligase, were purchased from New England Biolabs.

Nucleospin RNA extraction kit, GeneJET Gel Extraction Kit, GeneJET Plasmid Miniprep Kit, First-strand cDNA Synthesis Kit, 1 Kb Plus DNA Ladder, DreamTaq Green PCR Master Mix, Power SYBR Green PCR Master Mix and 0.2 μm micro-centrifugal filters were purchased from Thermo Fisher Scientific. GelRed nucleic acid gel stain was purchased from Biotium. Salmon sperm DNA was purchased form Agilent Technologies. Ninety-six-well microplates for RT-qPCR analysis were purchased from ABI Applied Biosystems. Primers for PCR and RT-qPCR amplification, as well as gBlocks, were purchased from Integrated DNA Technologies (IDT, see Supporting Table S2 for a list of oligonucleotides used in this study). Quick-DNA Fungal/Bacterial Miniprep Kit and Zymoprep Yeast Plasmid Miniprep I were purchased from Zymo Research. Drop-out mix minus histidine, leucine, lysine, methionine, uracil without yeast nitrogen base was purchased from US Biological. Coenzyme Q_6_ standard (*Saccharomyces cerevisiae*, >99% purity) was purchased from Sigma-Aldrich.

### Microorganisms and media

*S. cerevisiae* EPY300,^16^ a derivative of *S. cerevisiae* BY4742,^32^ was provided by Prof. Jay Keasling (UC Berkeley). This and derived strains EPYFA3, EPYFA4 and EPYFA7 were grown in synthetic defined (SD) broth (6.7 g L^−1^ yeast nitrogen base without amino acids, 1.318 g L^−1^ drop-out mix) with histidine, leucine, uracil and/or methionine dropped out as appropriate. Filter-sterilized dextrose and/or galactose were used as carbon sources and added after autoclaving. One Shot TOP10 chemically competent *E. coli* cells (Thermo Fisher Scientific) were used for cloning and storage of plasmid DNA. *E. coli* strains were grown on lysogeny broth (LB) medium (10 g L^−1^ tryptone, 5 g L^−1^ yeast extract, 10 g L^−1^ NaCl) or LB agar medium (same as LB medium, with 15 g L^−1^ agar), with appropriate antibiotic selection (25 μg mL^−1^ chloramphenicol, 100 μg mL^−1^ ampicillin, or 50 μg mL^−1^ kanamycin when transformed with pYTK-derived plasmids. The MoClo Yeast Toolkit^25^ was obtained from Addgene (Kit # 1000000061) and was a gift from Prof. John E. Dueber (UC Berkeley).

### Expression vector assembly

Plasmids for expression of *aroL*, *GCN4* and *COQ2* were assembled using the MoClo Yeast Toolkit^25^ according to the manufacturer’s instructions using Golden Gate assembly.^33^ The gene *aroL* was codon-optimized for expression in *S. cerevisiae*, obtained as a synthetic gBlock from IDT and cloned into pYTK001 to obtain the part plasmid pFA002, which was checked through Sanger sequencing. *GCN4* was amplified from the genomic DNA of *S. cerevisiae* EPY300 and cloned into pYTK001 to obtain the part plasmid pFA003. Cassette plasmids pFA007 and pFA008 were constructed for *aroL* and *GCN4*, respectively, using pFA002, pFA003 and the pre-assembled vector pYTK095 as a backbone. Finally, the multigene plasmid pFA011 was assembled to include the expression cassettes that comprised *aroL* and *GCN4* using as a backbone the GFP-drop out containing plasmid pFA010, which contained homologous recombination arms for integration in the *LEU2* locus, as well as the yeast selectable marker *LEU2*. Correct assembly of pFA011 was verified through restriction digestion (Supporting Figure S1) and Sanger sequencing. *COQ2* was obtained as a synthetic gBlock from IDT where a BsmbI site was removed through a synonymous substitution (G885A). We cloned *COQ2* into pYTK001, obtaining part plasmid pFA001, which was checked through Sanger sequencing and then used to produce the cassette plasmid pFA004 through Golden Gate, which included homologous recombination arms for integration in the *URA3* locus, as well as the yeast selectable marker *URA3*. Correct assembly of pFA004 was checked through restriction digestion (Supporting Figure S2) and Sanger sequencing. Golden Gate assembly reaction was performed as follows: 5.7 fmols of DNA vector were added with inserts in a 1:3 molar ratio (vector: insert), as well as with 1 µL of 10x T4 Ligase buffer, 0.5 µL of T4 Ligase, 0.5 µL of either 10,000 U mL^−1^ BsmBI (for part and multigene plasmid assembly) or 10,000 U mL^−1^ BsaI (for cassette plasmid assembly), then made up to a final volume of 10 µL with deionized water. The reaction mixture was incubated in a thermal cycler with the following program: 10 cycles of digestion and ligation (37°C for 10 minutes, 16°C for 10 minutes) followed by a final digestion step (50°C for 5 minutes), and a heat inactivation step (65°C for 20 minutes). The final digestion and heat inactivation steps were omitted, therefore ending on a ligation, for the assembly of the GFP-dropout containing plasmid pFA010.

## Author Contributions

D.K. and F.A. performed experiments. F.A. and C.C provided equipment and materials. F.A. conceived and designed the study and wrote the manuscript with the assistance of C.C.

All authors have given approval to the final version of the manuscript. The authors declare no conflict of interest.

## ACKNOWLEDGMENT

The authors would like to acknowledge E.M. Wellington for providing equipment. E. Silvano and Y. Chen are thanked for support and guidance with lipid extraction and analysis. J. Keasling (UC Berkeley) is thanked for providing strain *S. cerevisiae* EPY300. J.E. Dueber (UC Berkeley) is thanked for providing the MoClo Yeast Toolkit. The authors would like to acknowledge funding from the Leverhulme Trust (ECF-2018-691) and the University of Warwick, through an Early Career Fellowship to F.A., as well as from EPSRC and BBSRC (EP/L016494/1) for a scholarship to D.K. through the Centre for Doctoral Training in Synthetic Biology (SynBioCDT). The Warwick Integrative Synthetic Biology Centre (WISB), a UK Synthetic Biology Research Centre grant from EPSRC and BBSRC (BB/M017982/1), is also acknowledged.

