## Supporting Information for "Engineering isoprenoid quinone production in yeast"

### Supporting Methods

#### Strain engineering

A list of all strains used in this study is available in Supplementary Table S1. Strain EPY300 was used as a recipient host for the chromosomal integration of plasmid pFA011 (containing genes *aroL* and *GCN4*), to obtain strain EPYFA3, as well as of plasmid pFA004 (containing gene *COQ2*) to obtain strain EPYFA4, and of both plasmids pFA004 and pFA011 to obtain strain EPYFA7. Yeast transformation was performed according to the LiAc/single-stranded carrier DNA/PEG method.[1] Briefly, a single colony of EPY300 was picked up from a selective plate of SD agar (same as SD broth, with 15 g $L^{-1}$ agar) with histidine dropped out and used to inoculate five mL of SD broth with histidine dropped out, incubated overnight at 30°C, shaking at 200 rpm. The overnight culture was used to inoculate 50 mL of SD broth with histidine dropped out, incubated at 30°C, shaking at 200 rpm, until $OD_{600}$ = 0.7 was reached. Meanwhile a restriction digestion was set up for plasmids pFA004 and pFA011 as follows: 16 µL of plasmid DNA, 2 µL of Cutsmart buffer (NEB), 2 µL of NotI; the reaction was incubated at 37°C for 3 hours, after which 2 µL of rSAP (NEB) were added and the reaction incubated for one additional hour. The reaction was deactivated by incubating at 80°C for 20 minutes, then kept on ice. Once the appropriate cell density was reached, yeast cells were harvested by centrifugation and resuspended in 25 mL of sterile water, harvested again and resuspended in 0.1 M lithium acetate to a final volume of one mL. One hundred µL aliquots of yeast cells were prepared, harvested by centrifugation and supernatant removed. Each cell pellet was added with 360 µL of transformation mixture, comprising of 240 µL of 50% (w $v^{-1}$) PEG 3,350, 36 µL of 1.0 M lithium acetate, 10 µL of 10.0 mg $mL^{-1}$ boiled salmon sperm DNA, and 74 µL of plasmid DNA (pFA004 and/or pFA011) plus water. After vigorous vortexing, the transformation reaction was incubated at 30°C for 30 minutes, then at 42°C for 30 minutes. Cells were pelleted and resuspended in 500 µL of sterile water, then 200 µL of the cell suspension, as well as a 1:10 dilution of it, were spread onto SD agar selective plates with histidine, leucine and uracil dropped out as appropriate. Plates were incubated at 30°C for 3 days until colonies appeared. Colonies were streaked onto fresh SD agar selective plates with histidine, leucine and uracil dropped out as appropriate for downstream applications.

#### Nucleic acid purification, PCR and RT-qPCR

Genomic DNA was purified from *S. cerevisiae* strains EPY300, EPYFA3, EPYFA4 and EPYFA7 with the Quick-DNA Fungal/Bacterial Miniprep Kit (Zymo Research), following the manufacturer's instructions. Q5 High-Fidelity DNA polymerase (NEB) was used to amplify *GCN4* from the genomic DNA of *S. cerevisiae* EPY300. PCR was performed in a 25 µL volume as follows: 12.5 µL of Q5 High-Fidelity 2X Master Mix, 1.25 µL of each 10 µM forward and reverse primer, 1 µL of purified genomic DNA and 9 µL of deionized water. Thermal cycling was performed in a Bio-Rad T100 Thermal Cycler as follows: initial denaturation of 98°C for 2 minutes, followed by 35 cycles of DNA denaturation at 98°C for 10 seconds, primer annealing for 20 seconds, DNA amplification at 72°C for 30 seconds per kb amplified, and a final extension at 72°C for 5 minutes. The annealing temperature for specific primer pairs was calculated using the NEB Tm Calculator tool (http://tmcalculator.neb.com). The PCR product was purified from a 1% w $v^{-1}$ agarose gel using the GeneJET Gel Extraction Kit (Thermo Fisher Scientific), following the manufacturer's instructions, then used for downstream applications.

Purified genomic DNA from EPYFA3, EPYFA4, EPYFA7 was screened for correct chromosomal insertion of plasmids pFA011 and/or pFA004, using EPY300 as negative control. PCR was performed in a 25 µL volume with DreamTaq Green DNA Polymerase (Thermo Fisher Scientific), as follows: 12.5 µL of 2X DreamTaq Green PCR Master Mix, 1 µL of each 10 µM forward and reverse primer, 1 µL of purified genomic DNA and 9.5 µL of deionized water. Thermal cycling was performed in a Bio-Rad T100 Thermal Cycler as follows: initial denaturation of 95°C for 5 minutes, followed by 35 cycles of DNA denaturation at 95°C for 30 seconds, primer annealing at 50-65°C (calculated for specific primer pairs using the NEB Tm Calculator tool, http://tmcalculator.neb.com) for 30 seconds, DNA amplification at 72°C for 1 minute per kb amplified, and a final extension at 72°C for 5 minutes.

RNA was purified using the Nucleospin RNA extraction kit, including the optional step for in-column DNase I-digestion, according to the manufacturer's instructions; quality and quantity of RNA were assessed through gel electrophoresis (1% w $v^{-1}$ agarose gel with 1X GelRed) and NanoDrop ND1000 (Thermo Fisher Scientific). One µg of total RNA was used to set up a first-strand cDNA synthesis reaction using the First-strand cDNA Synthesis Kit (Thermo Fisher Scientific) with Oligo(dT)$_{18}$ primer, following the manufacturer's instructions. The resulting cDNA was quantified using NanoDrop ND1000 (Thermo Fisher Scientific) and normalized to a concentration of 50 ng µ$L^{-1}$. RT-qPCR was carried out to compare levels of expression of genes *ARO1*, *ARO2*, *ARO3*, *ARO4* and *GCN4* between strains EPYFA3 and EPY300, using *ACT1* as the internal control. Briefly, primers pairs for the amplification of each gene were designed using PrimeQuest (Integrated DNA Technologies, see Supporting Table S2). Three biological replicates (independent colonies) were tested for each of the two strains; three technical replicates were performed for each sample. RT-qPCR was performed in MicroAmp Optical 96-well reaction plates (Applied Biosystems) in a 20 µL volume with Power SYBR Green Master Mix (Thermo Fisher Scientific), as follows: 12.5 µL of 2X Power SYBR Green PCR Master Mix, 0.4 µL of each 10 µM forward and reverse primer, 1 µL of cDNA and 8.2 µL of deionized water. Thermal cycling was performed in a 7500HT fast real-time PCR system (Applied Biosystems) as follows: 50°C for 2 minutes and 95°C for 10 minutes for one cycle, followed by 40 cycles of DNA denaturation at 95°C for 15 seconds, primer annealing and extension at 60°C for 60 seconds; melting curve analysis was performed by heating to 95°C, followed by cooling to 60°C. The endogenous gene *ACT1* was used as an internal control to normalize the expression of the target genes.

**Lipid extraction and HPLC-MS analysis**

Lipids extraction was performed from yeast strains based on a modified protocol from Marbois *et al.*[2] Briefly, yeast strains were grown for 120 hours, $OD_{600}$ was recorded and used to collect cell cultures equivalent to 5 mL of an $OD_{600}$ = 1 culture. Cells were pelleted, supernatant discarded, and pellets were stored at -80°C. Cells were thawed on ice, 500 µL of ice-cold methanol were added and samples were vortexed for 30 seconds. Samples were added with 100 µL of water and 2 mL of chloroform, vortexed again and centrifuged at 3,000 rpm for 5 minutes at 4 °C with no brake. The chloroform phase was moved to a new glass vial, 2 mL of chloroform were added to the remaining water-methanol suspension, which was vortexed for 30 seconds and centrifuged as before. The chloroform phase was pooled with the previously collected organic phase and solvent was evaporated under nitrogen flow. The dried extracts were resuspended in 200 μL of 95% acetonitrile : 5% ammonium acetate (10 mM, pH 9.2), filtered through a 0.2 μm micro-centrifugal filter and moved to HPLC vials. For reversed-phase HPLC-MS analysis, an Agilent 1260 HPLC was used, equipped with Agilent 1260 Infinity diode array detector (DAD) VL, coupled to an AmaZon X quadrupole ion trap MS (Bruker) *via* an electrospray ionization interface, using a BEH Amide XP column (Waters) (2.5 μm, 3.0 x 150 mm XP) with a flow rate of 0.350 mL $min^{-1}$. UV DAD wavelength was set to 274 nm to detect coenzyme $Q_6$. A 1.1' delay between the DAD and the MS was consistently observed, due to the tubing between the devices, in both the authentic coenzyme $Q_6$ standard and all yeast crude extracts. Solvent A was acetonitrile, solvent B was water with 10 mM ammonium acetate, pH 9.3. The solvent gradient was as follows: 95:5 solvent A/solvent B for 10 minutes to equilibrate the column, followed by injection of 5 μL of sample, then 95:5 solvent A/solvent B to 70:30 solvent A/solvent B over 22 minutes.

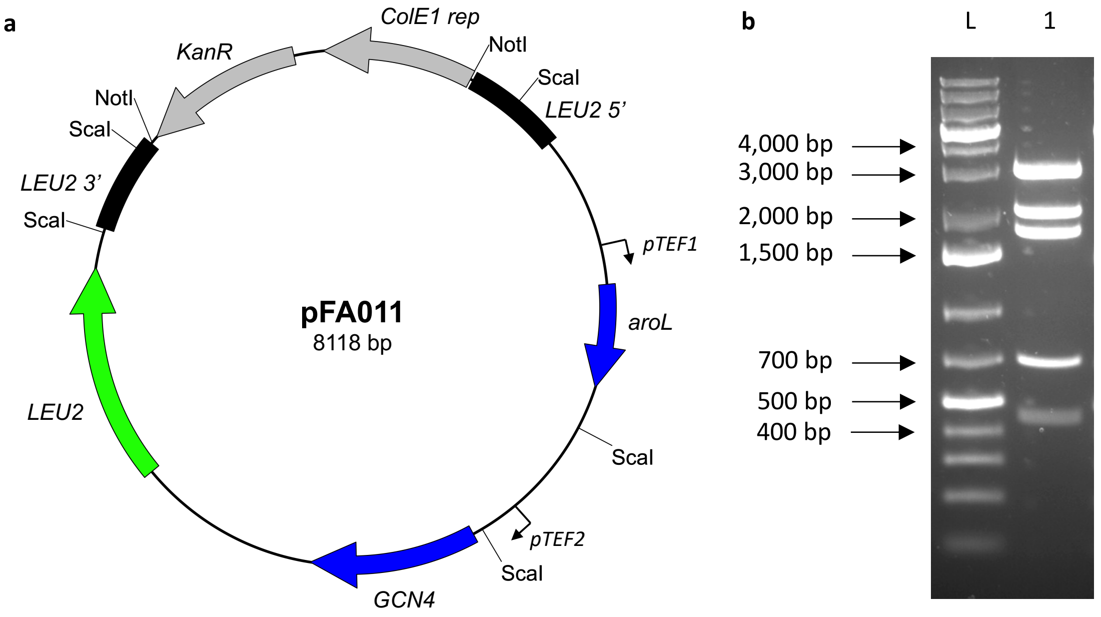

**Supporting Figure S1**. Assembly of plasmid pFA011. **(a)** Plasmid map. **(b)** L: 2 μL of GeneRuler 1kb Plus DNA Ladder (Thermo Scientific); lane 1: restriction digestion showing correct assembly of the plasmid. Expected bands upon digestion with ScaI: 3,104 bp; 2,090 bp; 1,769 bp; 707 bp; 448 bp.

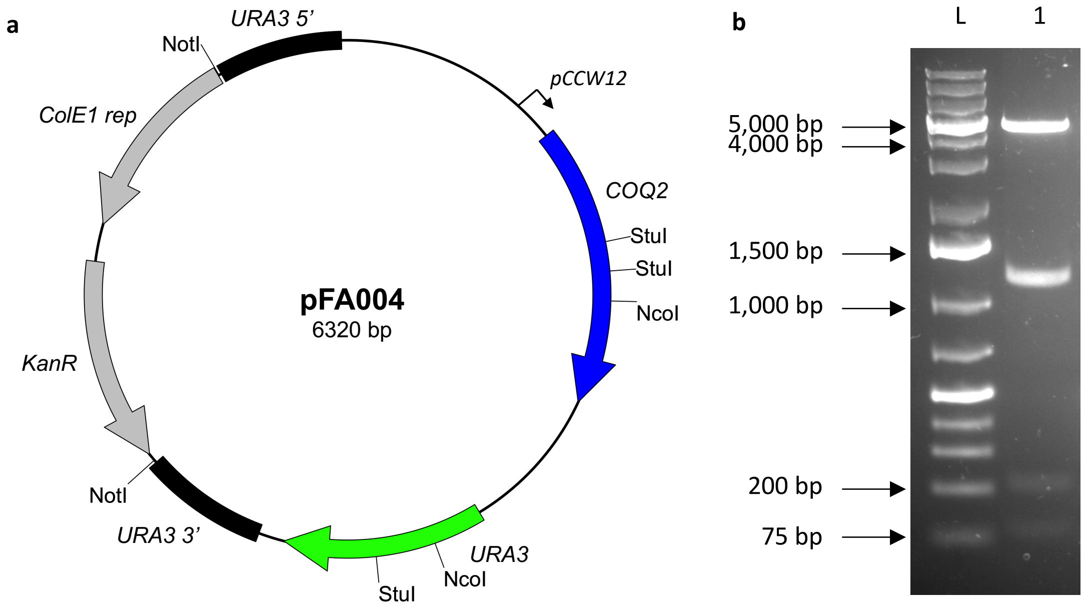

**Supporting Figure S2**. Assembly of plasmid pFA004. **(a)** Plasmid map. **(b)** L: 2 μL of GeneRuler 1kb Plus DNA Ladder (Thermo Scientific); lane 1: restriction digestion showing correct assembly of the plasmid. Expected bands upon digestion with NcoI and StuI: 4,649 bp; 1,213 bp; 231 bp; 114 bp.

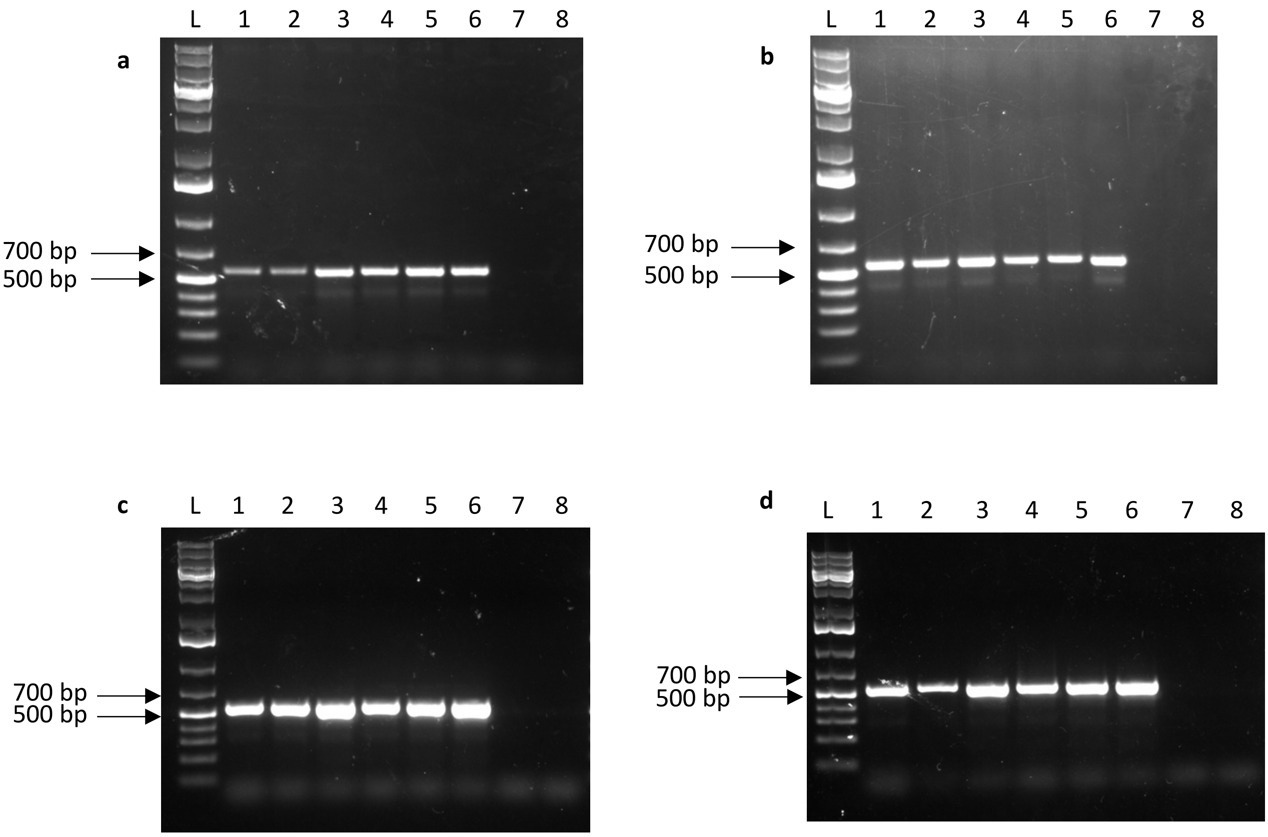

**Supporting Figure S3**. PCR screening for chromosomal integration in yeast strains. L: 2 µL of GeneRuler 1kb Plus DNA Ladder (Thermo Scientific). Correct integration into the target locus for strains EPYFA3, EPYFA4, EPYFA7 gives bands of ~ 540 bp, whereas the recipient strain (EPY300) gives no band.

**(a)** LEU2 5' integration screening on EPYFA3 colonies (lane 1-3), EPYFA7 colonies (lane 4-6), EPY300 (lane 7), negative control (lane 8). **(b)** LEU2 3' integration screening on EPYFA3 colonies (lane 1-3), EPYFA7 colonies (lane 4-6), EPY300 (lane 7), negative control (lane 8). **(c)** URA3 5' integration screening on EPYFA4 colonies (lane 1-3), EPYFA7 colonies (lane 4-6), EPY300 (lane 7), negative control (lane 8). **(d)** URA3 3' integration screening on EPYFA4 colonies (lane 1-3), EPYFA7 colonies (lane 4-6), EPY300 (lane 7), negative control (lane 8).

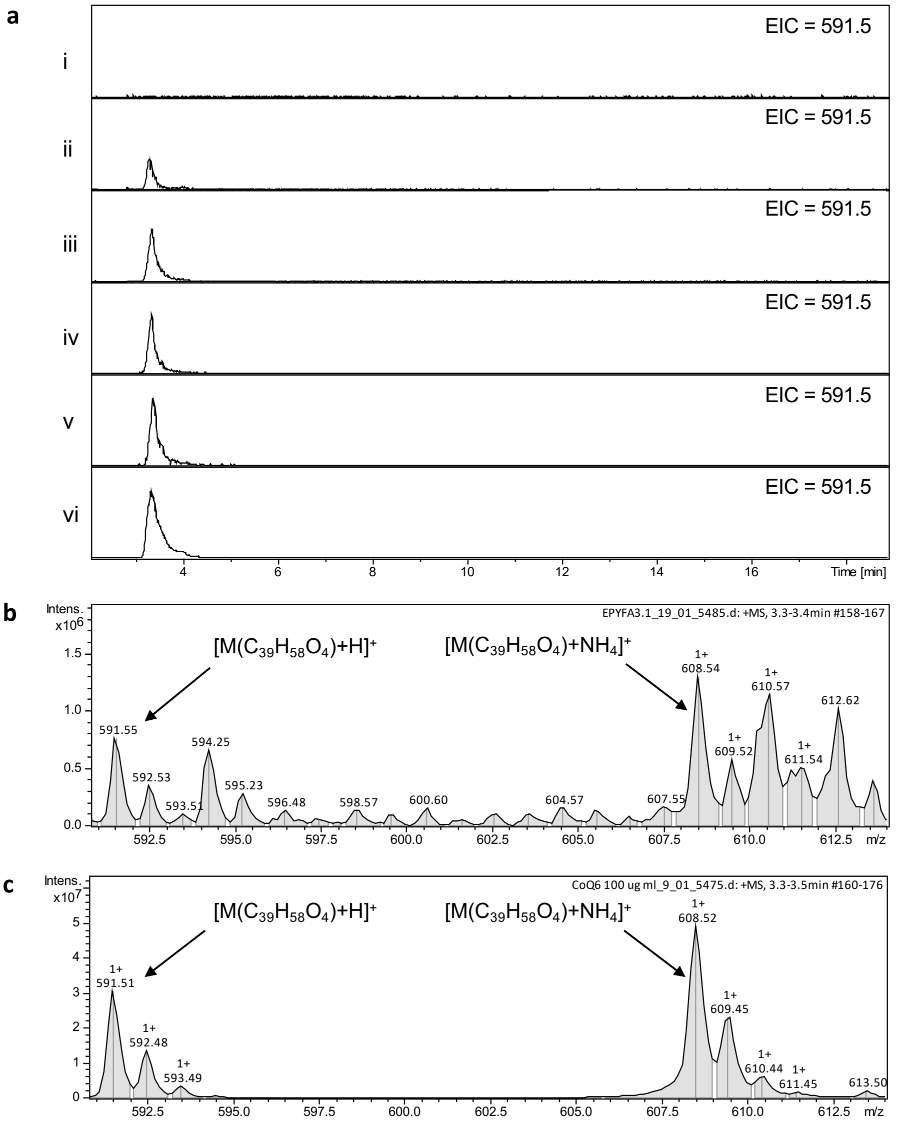

**Supporting Figure S4**. Mass spectrometry detection of coenzyme $Q_6$. **(a)** Extracted ion chromatograms in positive mode for *m/z* 591.5, corresponding to $[M(C_{39}H_{58}O_4)+H]^+$, are shown. Chromatograms from top to bottom: (i) growth medium control, (ii) EPY300, (iii) EPYFA3, (iv) EPYFA4, (v) EPYFA7, (vi) coenzyme $Q_6$ authentic standard. **(b)** mass spectrum of coenzyme $Q_6$ at retention time 3.3' from crude extract of EPY300. **(c)** mass spectrum of coenzyme $Q_6$ at retention time 3.3' for authentic standard.

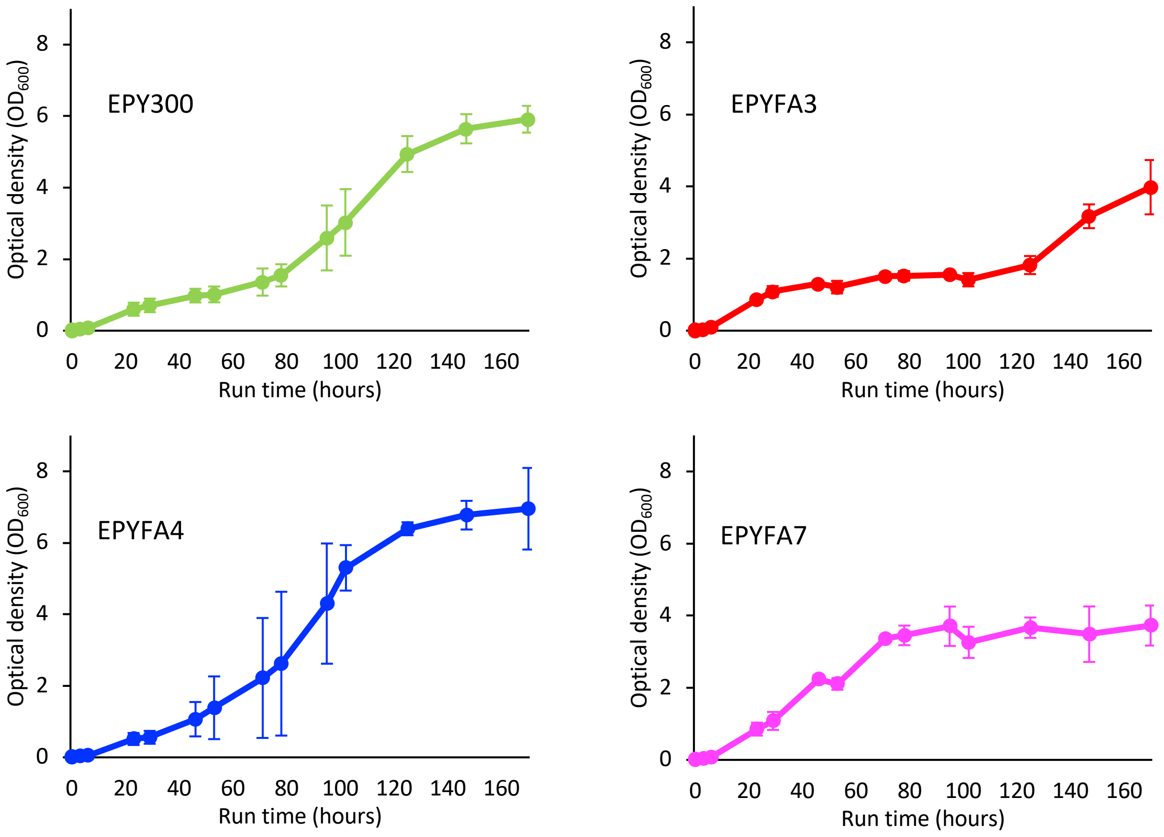

**Supporting Figure S5**. Growth of yeast strains in shake-flask cultures. EPY300 (green), EPYFA3 (red), EPYFA4 (blue), EPYFA7 (magenta).

**Supporting Table S1.** Strains and plasmids used in this study.

| Strain | Genotype | Source |
|---|---|---|
| *E. coli* One Shot TOP10 | F- *mcrA* Δ(*mrr-hsd*RMS-*mcr*BC) Φ80*lac*ZΔM15 Δ *lac*X74 *rec*A1 *ara*D139 Δ(*araleu*)7697 *gal*U *gal*K *rps*L (StrR) *end*A1 *nup*G | Thermo Fisher |
| *S. cerevisiae* EPY300 | $P_{GAL1}$-*tHMGR* $P_{GAL1}$-*upc2-1* *erg9::*$P_{MET3}$-*ERG9* $P_{GAL1}$-*tHMGR* $P_{GAL1}$-*ERG20* | Ro *et al.*[3] |
| *S. cerevisiae* EPYFA3 | $P_{GAL1}$-*tHMGR* $P_{GAL1}$-*upc2-1* *erg9::*$P_{MET3}$-*ERG9* $P_{GAL1}$-*tHMGR* $P_{GAL1}$-*ERG20* $P_{TEF1}$-*aroL* $P_{TEF2}$-*GCN4* | This work |
| *S. cerevisiae* EPYFA4 | $P_{GAL1}$-*tHMGR* $P_{GAL1}$-*upc2-1* *erg9::*$P_{MET3}$-*ERG9* $P_{GAL1}$-*tHMGR* $P_{GAL1}$-*ERG20* $P_{CCW12}$-*COQ2* | This work |
| *S. cerevisiae* EPYFA7 | $P_{GAL1}$-*tHMGR* $P_{GAL1}$-*upc2-1* *erg9::*$P_{MET3}$-*ERG9* $P_{GAL1}$-*tHMGR* $P_{GAL1}$-*ERG20* $P_{TEF1}$-*aroL* $P_{TEF2}$-*GCN4* $P_{CCW12}$-*COQ2* | This work |
| Plasmid | Content | Source |
| pFA001 | *COQ2 CamR* ColE1 | This work |
| pFA002 | *aroL CamR* ColE1 | This work |
| pFA003 | *GCN4 CamR* ColE1 | This work |
| pFA004 | ConLS $P_{CCW12}$-*COQ2*-$t_{ENO}$ ConRE *URA3* URA3_3'_Hom *KanR*-ColE1 URA3_5’_Hom | This work |
| pFA007 | ConLS $P_{TEF1}$-*aroL*-$t_{SSA1}$ ConR1 *AmpR*-ColE1 | This work |
| pFA008 | ConL1 $P_{TEF2}$-*GCN4*-$t_{ADH1}$ ConRE *AmpR*-ColE1 | This work |
| pFA010 | ConLS’ $P_{BBa\_J72163\ GlpT}$-*sfGFP*-$t_{BBa\_B0015}$ ConRE’ *LEU2* LEU2_3’_Hom *KanR*-ColE1 LEU2_5’_Hom | This work |
| pFA011 | ConLS $P_{TEF1}$-*aroL*-$t_{SSA1}$ ConR1 ConL1 $P_{TEF2}$-*GCN4*-$t_{ADH1}$ ConRE *LEU2* LEU2_3’_Hom *KanR*-ColE1 LEU2_5’_Hom | This work |

**Supporting Table S2**. List of oligonucleotides used in this work.

| Name | Sequence (5' → 3') | Description |
|---|---|---|
| *aroL* | ATGACCCAACCTTTGTTTTTGATTGGTCCAAGAGGTTGTGGTAAGACTACTGTTGGTATGGCTTTGGCTGATTCTTTGAACAGAAGATTCGTTGATACCGACCAATGGTTGCAATCTCAATTGAATATGACCGTTGCCGAAATCGTTGAAAGGGAAGAATGGGCTGGTTTTAGAGCTAGAGAAACTGCTGCTTTGGAAGCTGTTACTGCTCCATCTACTGTTATTGCTACTGGTGGTGGTATTATCTTGACCGAATTCAACAGACACTTCATGCAAAACAACGGTATCGTTGTCTATTTGTGTGCTCCAGTTTCTGTTTTGGTCAACAGATTGCAAGCTGCTCCAGAAGAAGATCTAAGACCAACTTTGACTGGTAAGCCATTGTCTGAAGAAGTTCAAGAAGTCTTGGAAGAAAGAGATGCCTTGTATAGAGAAGTTGCCCATATTATCATTGACGCTACCAATGAACCATCTCAGGTTATCTCTGAAATTAGATCTGCTTTGGCCCAAACCATTAACTGCTAA | *E. coli* shikimate kinase II gene, codon-optimized for expression in *S. cerevisiae* |
| FA001 | GCATCGTCTCATCGGTCTCATATGTCCGAATATCAGCCAAG | Amplification of *GCN4* from yeast genomic DNA and assembly into plasmid pFA003 |
| FA002 | ATGCCGTCTCAGGTCTCAGGATTCAGCGTTCGCCAACTAAT | |
| FA003 | GGTTCCTGGCCTTTTGCTGG | Sequencing of plasmids pFA003, pFA004, pFA007, pFA008, pFA011 |
| FA004 | CCTGTTGATAGATCCAGTAA | Sequencing of plasmid pFA003 |
| FA005 | GCCCACATGGATAACATTA | Sequencing of plasmid pFA004 |
| FA006 | CATGGCCTTCATGATAGAC | Sequencing of plasmid pFA004 |
| FA007 | ACTATATACGCACACCAGG | Sequencing of plasmid pFA004 |
| FA008 | GCAGTGGATTCAAGTGCTC | Sequencing of plasmid pFA004 |
| FA009 | TTGAGAAGATGCGGCCAGC | Sequencing of plasmid pFA004 |
| FA010 | CATCACGGGCAAGGATTGT | Sequencing of plasmid pFA0011 |
| FA011 | TGTACTGCAGTGCACTAGT | Sequencing of plasmids pFA008, pFA0011 |
| FA012 | GGCGACTCACAACTAAGCA | Sequencing of plasmid pFA008 |
| FA013 | ATGTAGATTGCGTATATAG | Sequencing of plasmid pFA0011 |
| FA014 | CTCGTATCGCATGTCGGTG | Sequencing of plasmid pFA0011 |
| FA015 | ATGATGAGCCGTGATGACC | Sequencing of plasmid pFA007 |
| FA016 | CTTCGACTATGCTGGAGG | Sequencing of plasmid pFA011 |
| FA017 | ATATGTAAGTATACGGCC | Sequencing of plasmid pFA011 |
| FA018 | GGGCGGATTACTACCGTT | Amplification of URA3 5' region for verification of chromosomal integrations |
| FA019 | GTAATGTTATCCATGTGGGC | |
| FA020 | AGAGCACTTGAATCCACTGC | Amplification of URA3 3' region for verification of chromosomal integrations |
| FA021 | GATTTGGTTAGATTAGATATGGTTTC | |
| FA022 | CATAAATACCTTTCAAGC | Amplification of LEU2 5' region for verification of chromosomal integrations |
| FA023 | TACAATCCTTGCCCGTGATG | |
| FA024 | ACTCGTATCGCATGTCGGTG | Amplification of LEU2 3' region for verification of chromosomal integrations |
| FA025 | CTTCTTATGTTTTACATG | |
| FA133 | GGGTCCTCCTGTAGGTACTTTA | Amplification of *ARO1* for qPCR analysis |
| FA134 | GGATCACTGTCGTGCGAAATA | |

| | | |
|---|---|---|
| FA135 | CAGAGGCAACAAGGACTCTATC | Amplification of *ARO2* for qPCR analysis |
| FA136 | CAACATGGCTTCCAACTTGTC | |
| FA137 | TTCAGCCAAAGGTGAGGAAA | Amplification of *ARO3* for qPCR analysis |
| FA138 | GGCCCGATCACGATAACTAAA | |
| FA139 | GAAACTGCCAAGAGAGGTAGAA | Amplification of *ARO4* for qPCR analysis |
| FA140 | GGAACAAGGACCGACAATGA | |
| FA141 | ACGCTGTAGTGGAATCTTTCTT | Amplification of *GCN4* for qPCR analysis |
| FA142 | CGTCAGTGGTAACTGGAATGT | |
| FA143 | CGTCTGGATTGGTGGTTCTATC | Amplification of *ACT1* for qPCR analysis |
| FA144 | GGACCACTTTCGTCGTATTCTT | |
